# Density-dependent selection and the limits of relative fitness

**DOI:** 10.1101/102087

**Authors:** Jason Bertram, Joanna Masel

## Abstract

Selection is commonly described by assigning constant relative fitness values to genotypes. Yet population density is often regulated by crowding. Relative fitness may then depend on density, and selection can change density when it acts on a density-regulating trait. When strong density-dependent selection acts on a density-regulating trait, selection is no longer describable by density-independent relative fitnesses, even in demographically stable populations. These conditions are met in most previous models of density-dependent selection (e.g. “*K*-selection” in the logistic and Lotka-Volterra models), suggesting that density-independent relative fitnesses must be replaced with more ecologically explicit absolute fitnesses unless selection is weak. Here we show that density-independent relative fitnesses can also accurately describe strong density-dependent selection under some conditions. We develop a novel model of density-regulated population growth with three ecologically intuitive traits: fecundity, mortality, and competitive ability. Our model, unlike the logistic or Lotka-Volterra, incorporates a density-dependent juvenile “reproductive excess”, which largely decouples density-dependent selection from the regulation of density. We find that density-independent relative fitnesses accurately describe strong selection acting on any one trait, even fecundity, which is both density-regulating and subject to density-dependent selection. Pleiotropic interactions between these traits recovers the familiar *K*-selection behavior. In such cases, or when the population is maintained far from demographic equilibrium, our model offers a possible alternative to relative fitness.

## Introduction

There are a variety of different measures of fitness, such as expected lifetime reproductive ratio *R*_0_, intrinsic population growth rate *r*, equilibrium population density/carrying capacity (often labeled “*K*”) (Benton and Grant, 2000), and invasion fitness (Metz et al., 1992). In addition, “relative fitness” is widely used in evolutionary genetics, where the focus is on relative genotypic frequencies (Barton et al., 2007, pp. 468). The justification of any measure of fitness ultimately derives from how it is connected to the processes of birth and death which drive selection (Metcalf and Pavard 2007; Doebeli et al. 2017; Charlesworth 1994, pp. 178). While such a connection is clear for absolute fitness measures like *r* or *R*_0_, relative fitness has only weak justification from population ecology. It has even been proposed that relative fitness be justified from measure theory, abandoning population biology altogether (Wagner, 2010). Given the widespread use of relative fitness in evolutionary genetics, it is important to understand its population ecological basis, both to clarify its domain of applicability, and as part of the broader challenge of synthesizing ecology and evolution.

For haploids tracked in discrete time, the change in the abundance *n*_*i*_ of type *i* over a time step can be expressed as Δ*n*_*i*_ = (*W*_*i*_ − 1)*n*_*i*_ where *W*_*i*_ is “absolute fitness” (i.e. the abundance after one time step is 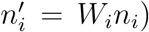. The corresponding change in frequency is.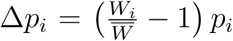 where 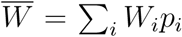. In continuous time, the Malthusian parameter *r*_*i*_replaces *W*_*i*_ and we have 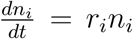 and 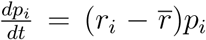 (Crow et al., 1970). Note that we can replace the *W*_*i*_ with any set of values proportional to the *W*_*i*_ without affecting the ratio 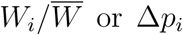 or Δ*p*_*i*_. These “relative fitness” values tell us how type frequencies change, but give no information about the dynamics of total population density *N* =Σ_*i*_ *n*_*i*_ (Barton et al., 2007, pp. 468). Similarly in the continuous case, adding an arbitrary constant to the Malthusian parameters *r*_*i*_ has no effect on 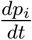 (these would then be relative log fitnesses).

Relative fitness is often parameterized in terms of selection coefficients which represent the advantages of different types relative to each other. For instance, in continuous time *s* = *r*_2_ *-r*_1_ is the selection coefficient of type 2 relative to type 1. Assuming that only 2 and 1 are present, the change in frequency can be written as

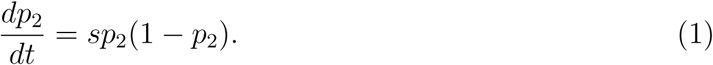

Thus, if *r*_1_ and *r*_2_ are constant, the frequency of the second type will grow logistically with a constant rate parameter *s*. We then say that selection is independent of frequency and density. The discrete time case is more complicated. Defining the selection coefficient by *W*_2_ = (1 + *s*)*W*_1_, and again assuming 1 and 2 are the only types present, we have

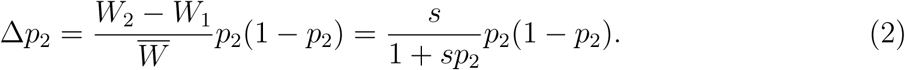

We will refer to both the continuous and discrete time selection equations (1) and (2) throughout this paper, but the simpler continuous time case will be our point of comparison for the rest of this section.

In a constant environment, and in the absence of crowding, *r*_*i*_ is a constant “intrinsic” population growth rate. The interpretation of Eq. (1) is then simple: the selection coefficient *s* is simply the difference in intrinsic growth rates. However, growth cannot continue at a non-zero constant rate indefinitely: the population is not viable if *r*_*i*_ *<* 0, whereas *r*_*i*_ *=* 0 implies endlessly increasing population density. Thus, setting aside unviable populations, the increase in population density must be checked by crowding. This implies that the Malthusian parameters *r*_*i*_ eventually decline to zero (e.g. Begon et al. 1990, pp. 203). Selection can then be density-dependent, and indeed this is probably not uncommon, because crowded and uncrowded conditions can favor very different traits (Travis et al., 2013). Eq. (1) is then not a complete description of selection — it lacks an additional coupled equation describing the dynamics of *N*, on which *s* in Eq. (1) now depends. In general we cannot simply specify the dynamics of *N* independently, because those ecological dynamics are coupled with the evolutionary dynamics of type frequency (Travis et al., 2013). Thus, in the presence of density-dependent selection, the simple procedure of assigning constant relative fitness values to different types has to be replaced with an ecological description of absolute growth rates. Note that frequency-dependent selection does not raise a similar problem, because a complete description of selection still only requires us to model the type frequencies, not the ecological variable *N* as well.

In practice, many population genetic models simply ignore density dependence and assign a constant relative fitness to each type. Selection is typically interpreted as operating through viability, but the ecological processes underlying the regulation of population density are frequently left unspecified (e.g. Gillespie 2010; Nagylaki et al. 1992; Ewens 2004). Density either does not enter the model at all, or if finite-population size effects (”random genetic drift”) are important, then *N* is typically assumed to have reached some fixed equilibrium value (Fig. 1b; for some approaches to relaxing the constant *N* assumption in finite populations, see Lambert et al. 2005; Parsons and Quince 2007; Chotibut and Nelson 2017; Constable and McKane 2017).

**Figure 1:**
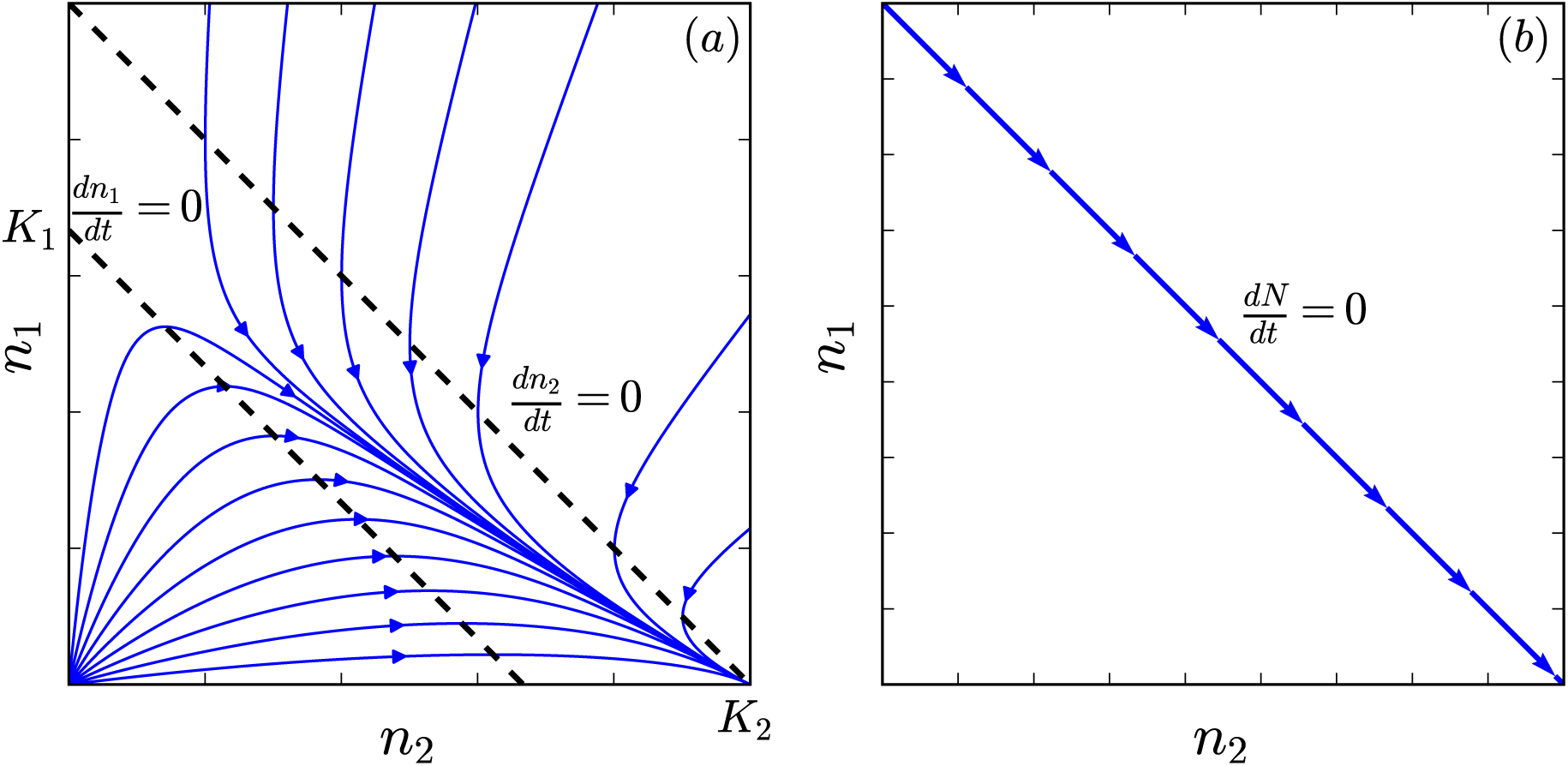
Phase diagram for the densities of two types *n*_1_ and *n*_2_ undergoing selection. (a) The logistic model 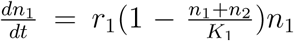 and 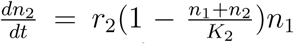 with *r*_1_ = *r*_2_ and *K*_2_ *> K*_1_. (b) The constant-*N*, relative fitness description of selection.

A rather different picture emerges in more ecologically explicit studies of selection in density-regulated populations. Following Fisher’s suggestion that evolution tends to increase density in the long term (Fisher, 1930; Leon and Charlesworth, 1978; Lande et al., 2009), as well as the influential concept of *K*-selection (specifically, the idea that selection in crowded conditions favors greater equilibrium density; MacArthur 1962), many studies of density-regulated growth have focused on the response of density to selection (Kostitzin, 1939; MacArthur and Wilson, 1967; Roughgarden, 1979; Christiansen, 2004). Indeed, both *N* and *s* change during, and as a result of, adaptive sweeps in many of the most widely used models of density-regulated population growth. The latter includes simple birth-death (Kostitzin, 1939) and logistic models (Fig. 1a; MacArthur 1962; Roughgarden 1979; Boyce 1984), variants of these models using other functional forms for the absolute fitness penalties of crowding (Kimura, 1978; Charlesworth, 1971; Lande et al., 2009; Nagylaki, 1979; Lande et al., 2009), and the “*R\** rule” of resource competition theory (which states that the type able to deplete a shared limiting consumable resource to the lowest equilibrium density *R\** excludes the others; Grover 1997). Density also changes in response to selection in the Lotka-Volterra competition model, at least during a sweep (except in special cases; Gill 1974; Smouse 1976; Mallet 2012).

The constant-*N*, constant-*s* description of selection also limits consideration of longer-term aspects of the interplay between evolution and ecology such as population extinction and trait evolution. A variety of approaches have been developed to address this in quantitative genetics (Burger and Lynch, 1995; Engen et al., 2013), population genetics (Bertram et al., 2017) and adaptive dynamics (Ferriére and Legendre, 2013; Dieckmann and Ferrière, 2004). Although density-dependent selection is pertinent to these longer-term issues, our focus here is the description of the time-dependent process by which selection changes allele frequencies. This is particularly critical for making sense of evolution at the genetic level, for which we now have abundant data.

In light of the complications arising from density-dependence, the assignment of density-independent relative fitnesses has been justified as an approximation that holds when selection is weak and *N* changes slowly (Kimura and Crow 1969; Ewens 2004, pp. 277; Charlesworth 1994, Chap. 4). Under these conditions, *s* is approximately constant in Eq. (1), at least for some number of generations. If *s* depends only on density, not frequency, this approximate constancy can hold over entire selective sweeps (Otto and Day, 2011).

However, the preceding arguments do not imply that the constant relative fitness idealization of population genetics *only* applies when selection is weak and *N* is stable (or when selection is actually density-independent). The idealization of assigning relative fitness values to genotypes is powerful, and so it is important to understand the specifics of when and how it succeeds or fails when selection is not weak, or *N* is not stable. For instance, in wild *Drosophila*, strong seasonally-alternating selection happens concurrently with large “boom-bust” density cycles (Messer et al., 2016; Bergland et al., 2014). Are we compelled to switch to a more ecologically-detailed model of selection based on Malthusian parameters or birth/death rates in this important model system? And if we make this switch, how much ecological detail do we need?

Here we argue that the simplified models of density-regulated growth mentioned above are potentially misleading in their representation of the interplay between selection and density. This ultimately derives from their failure to account for “reproductive excess”, that is, an excess of juveniles that experience stronger selection than their adult counterparts (Turner and Williamson, 1968). By allowing selection to be concentrated at a juvenile “bottleneck”, reproductive excess makes it possible for the density of adults to remain constant even under strong selection. Reproductive excess featured prominently in early debates about the regulation of population density (e.g. Nicholson 1954), and also has a long history in evolutionary theory, particularly related to Haldane’s “cost of selection” (Haldane, 1957; Turner and Williamson, 1968). Additionally, reproductive excess is implicit in foundational evolutionary-genetic models like the Wright-Fisher, where each generation involves the production of an infinite number of zygotes, of which a constant number *N* are sampled to form the next generation of adults. Likewise in the Moran model, a juvenile is always available to replace a dead adult every iteration no matter how rapidly adults are dying, and as a result *N* remains constant.

Nevertheless, studies of density-dependent selection rarely incorporate reproductive excess. This requires that we model a finite, density-dependent excess, which is substantially more complicated than modeling either zero (e.g. logistic) or infinite (e.g. Wright-Fisher) reproductive excess. Nei’s “competitive selection” model incorporated a finite reproductive excess to help clarify the “cost of selection” (Nei, 1971; Nagylaki et al., 1992), but used an unusual representation of competition based on pairwise interactions defined for at most two different genotypes, and was also restricted to equal fertilities for each genotype.

In models with detailed age structure, it is often assumed that the density of a “critical age group” mediates the population’s response to crowding (Charlesworth, 1994, pp. 54). Reproductive excess is a special case corresponding to a critical pre-reproductive age group. A central result of the theory of density-regulated age-structured populations is that selection proceeds in the direction of increasing equilibrium density in the critical age group (Charlesworth, 1994, pp. 148). This is a form of the classical *K*-selection ideas discussed above, but restricted to the critical age group (juveniles, in this case). The interdependence of pre-reproductive selection and reproductive density is thus overlooked as a result of focusing on density in the critical age group.

We re-evaluate the validity of the constant relative fitness description of selection in a novel model of density-regulated population growth that has a finite reproductive excess.

Our model is inspired by the classic discrete-time lottery model, which was developed by ecologists to study competition driven by territorial contests in reef fishes and plants (Sale, 1977; Chesson and Warner, 1981), and which has some similarities to the Wright-Fisher model (Svardal et al., 2015). Each type is assumed to have three traits: fecundity *b*, mortality *d*, and competitive ability *c*. In each iteration of the classic lottery model, each type produces a large number of juveniles, such that *N* remains constant (infinite reproductive excess). Competitive ability *c* affects the probability of winning a territory, and behaves like a pure relative fitness trait. Thus, fitness involves a product of fertility and juvenile viability akin to standard population genetic models of selection (e.g. Crow et al. 1970, pp. 185). We relax the large-juvenile-number assumption of the lottery model to derive a variable-density lottery with a finite, density-dependent reproductive excess.

The properties of density-dependent selection in our model are strikingly different from the classical literature discussed above. The strong connection between crowding and selection for greater equilibrium density is broken: selection need not affect density at all. And when it does, the density-independent discrete-time selection equation (2) is almost exact even for strong selection, provided that any changes in density are driven only by selection (as opposed to large deviations from demographic equilibrium), and that selection occurs on only one of the traits *b, c*, or *d*. On the flip side, the constant relative fitness approximation fails when strong selection acts concurrently on two or more of these traits, or when the population is far from demographic equilibrium.

## Model

### Assumptions and definitions

We restrict our attention to asexual haploids, since it is then clearer how the properties of selection are tied to the underlying population ecological assumptions. We assume that reproductively mature individuals (”adults”) require their own territory to survive and reproduce. All territories are identical, and the total number of territories is *T.* Time advances in discrete iterations, each representing the time from birth to reproductive maturity. In a given iteration, the number of adults of the *i*’th type will be denoted by *n*_*i*_, the total number of adults by *N* = Σ_*i*_ *n*_*i*_, and the number of unoccupied territories by *U* = *T -N.* We assume that the *n*_*i*_ are large enough that stochastic fluctuations in the *n*_*i*_ (drift) can be ignored, with *T* also assumed large to allow for low type densities *n*_*i*_*/T ≪* 1.

Each iteration, adults produce propagules which disperse at random, independently of distance from their parents, and independently of each other (undirected dispersal). We assume that each adult from type *i* produces *b*_*i*_ propagules on average, so that the mean number of *i* propagules dispersing to unoccupied territories is *m*_*i*_ = *b*_*i*_*n*_*i*_*U/T* (the factor *U/T* represents the loss of those propagules landing on occupied territories). Random dispersal is modeled using a Poisson distribution 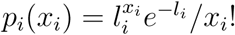 for the number *x*_*i*_ of *i* propagules dispersing to any particular unoccupied territory, where *l*_*i*_ = *m*_*i*_*/U* is the mean propagule density of type *i* per unoccupied territory. The total propagule density per unoccupied territory will be denoted *L* = _*i*_ *l*_*i*_.

We assume that adults cannot be ousted by juveniles, so that recruitment to adulthood occurs exclusively in unoccupied territories. When multiple propagules land on the same unoccupied territory, the winner is determined by lottery competition: type *i* wins a territory with probability *c*_*i*_*x*_*i*_*/*Σ _*i*_ *c*_*i*_*x*_*i*_, where *c*_*i*_ is a constant representing relative competitive ability (Fig. 2). Since the expected fraction of unoccupied territories with propagule composition *x*_1_*,…, x*_*G*_ is *p*_1_(*x*_1_) *p*_*G*_(*x*_*G*_) where *G* is the number of types present, and type *i* is expected to win a proportion *c*_*i*_*x*_*i*_*/*Σ _*i*_*c*_*i*_*x*_*i*_ of these, type *i*’s expected territorial acquisition is given by

**Figure 2:**
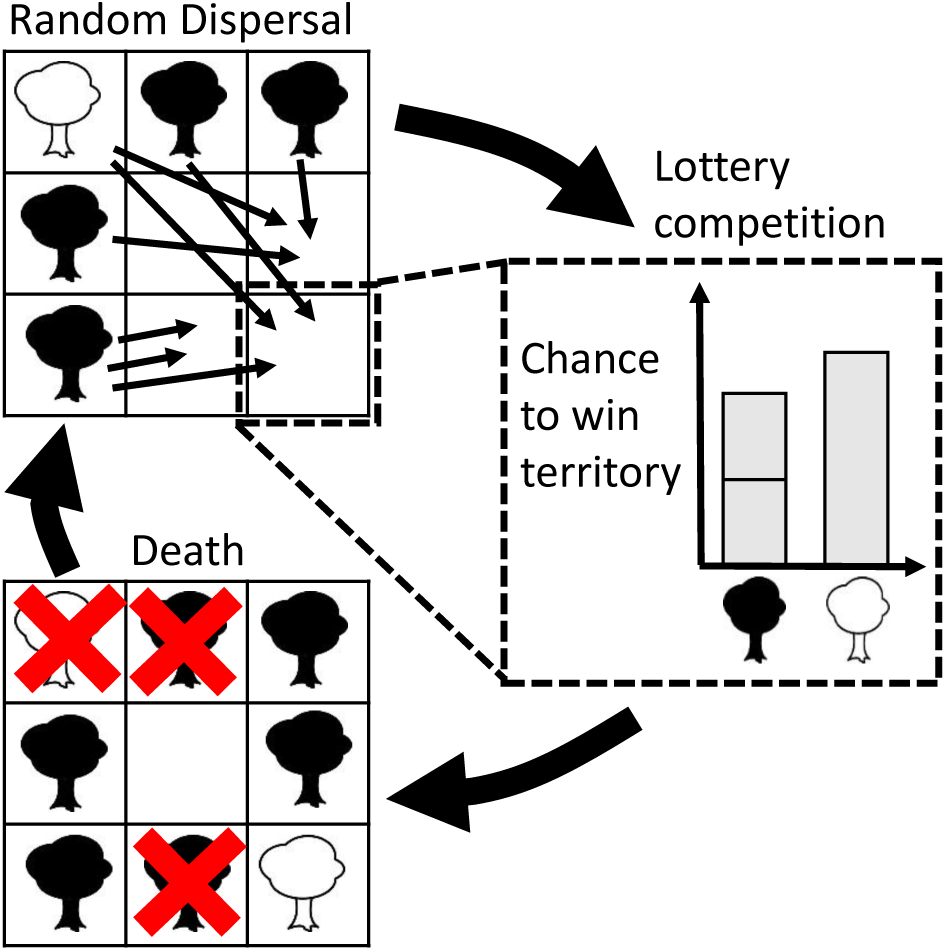
One iteration of our model. Propagules are dispersed by adults at random (only those propagules landing on unoccupied territories are shown). Some territories may receive zero propagules. Lottery competition then occurs in each territory that receives more than one propagule (only illustrated in one territory). In a given territory, type *i* has probability proportional to *c*_*i*_*x*_*i*_ of winning the territory, where *c*_*i*_ measures competitive ability and *x*_*i*_ is the number of *i* propagules present. In the illustrated territory, more black propagules are present, but white is a stronger competitor and has a higher probability of winning. Adult deaths make new territories available for the next iteration (red crosses).

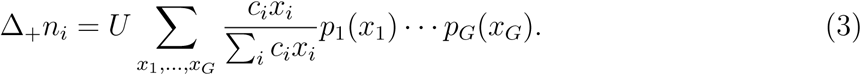

Here the sum only includes territories with at least one propagule present. Note that Δ_+_*n*_*i*_ denotes the expected territorial acquisition. Fluctuations about Δ_+_*n*_*i*_ (i.e. drift) will not be analyzed in this manuscript. Note that drift can become important if *U* is not sufficiently large even though *n*_*i*_ and *T* are large (by assumption); we do not consider this scenario on biological grounds, since it implies negligible population turnover.

Adult mortality occurs after lottery recruitment at a constant, type-specific per-capita rate *d*_*i*_ *≥* 1, and can affect adults recruited in the current iteration, such that the new abundance at the end of the iteration is (*n*_*i*_ + Δ_+_*n*_*i*_)*/d*_*i*_ (Fig. 2). In terms of absolute fitness, this can be written as

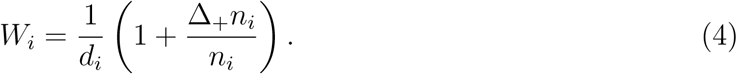

Here 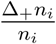 is the per-capita rate of territorial acquisition, and 1*/d*_*i*_ is the fraction of type *i* adults surviving to the next iteration. Note that our model of mortality differs from the classic lottery model (Chesson and Warner, 1981), where mortality affects adults only and occurs after propagule production but before juvenile recruitment. In the latter case, selection on mortality exhibits some density-dependence, although this reflects the fact that newly recruited adults are guaranteed to reproduce before dying, which is not interesting for our purposes here. Our mortality model ensures that selection on *d*_*i*_ is density-independent, allowing us to more clearly separate different sources of density-dependence and density regulation.

### Connection to the classic lottery model

In the classic lottery model (Chesson and Warner, 1981), unoccupied territories are assumed to be saturated with propagules from every type (*l*_*i*_ *→ ∞* for all *i*). From the law of large numbers, the composition of propagules in each territory will not deviate appreciably from the mean composition *l*_1_*, l*_2_*,…, l*_*G*_. Type *i* is thus expected to win a proportion *c*_*i*_*l*_*i*_*/Σ* _*i*_ *c*_*i*_*l*_*i*_ of the *U* available territories,

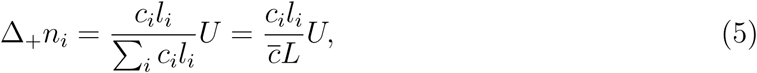

where 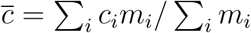 is the mean competitive ability for a randomly selected propagule. Note that all unoccupied territories are filled in a single iteration of the classic lottery model, whereas our more general model Eq. (3) allows for territories to be left unoccupied and hence also accommodates low propagule densities.

## Results

### Analytical approximation of the variable-density lottery

Here we evaluate the expectation in Eq. (3) to better understand the dynamics of density-dependent lottery competition. Similarly to the classic lottery model, we replace the *x*_*i*_, which take different values in different territories, with “effective” mean values. However, since we want to allow for low propagule densities, we cannot simply replace the *x*_*i*_ with the means *l*_*i*_ as in the classic lottery. For a low density type, growth comes almost entirely from territories with *x*_*i*_ = 1, for which its mean density *l*_*i*_ *⟪* 1 is not representative. We therefore separate Eq. (3) into *x*_*i*_ = 1 and *x*_*i*_ *>* 1 components, taking care to ensure that the effective mean approximations for these components are consistent with each other (details in Appendix A). The resulting variable-density approximation only requires that there are no large discrepancies in competitive ability (i.e. we do not have *c*_*i*_*/c*_*j*_ *≫* 1 for any two types). We obtain

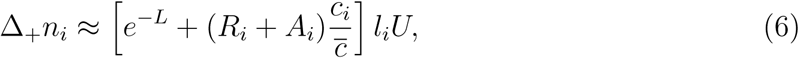

Where

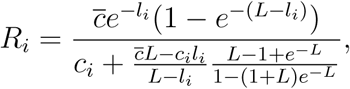

And

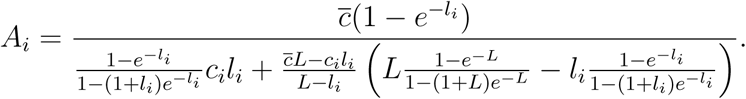

Comparing Eq. (6) to Eq. (5), the classic lottery per-propagule success rate 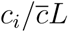 has been replaced by three separate terms. The first, *e*^*-L*^, accounts for propagules which land alone on unoccupied territories; these propagules secure the territories without contest. The second, 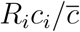 represents competitive victories on territories where only a single *i* propagule lands, together with at least one other propagule from a different type (this term dominates the growth of a rare invader in a high density population and determines invasion fitness). The third term, 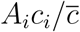 represents competitive victories in territories where two or more *i* type propagules are present. The relative importance of these three terms varies with both the overall propagule density *L* and the relative propagule frequencies *l*_*i*_*/L*. If *l*_*i*_ *≫* 1 for all types, we recover the classic lottery model (only the 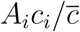 term remains, and *A*_*i*_ *→* 1*/L*).

Fig. 3 shows that Eq. (6) and its components closely approximate simulations of our variable-density lottery model over a wide range of propagule densities. Two types are present, one of which is at low frequency. The growth of the low-frequency type relies crucially on the low-density competition term.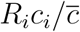 On the other hand, 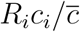 is negligible for the high-frequency type, which depends instead on high density territorial victories. Fig. 3 also shows the breakdown of the classic lottery model at low propagule densities.

**Figure 3:**
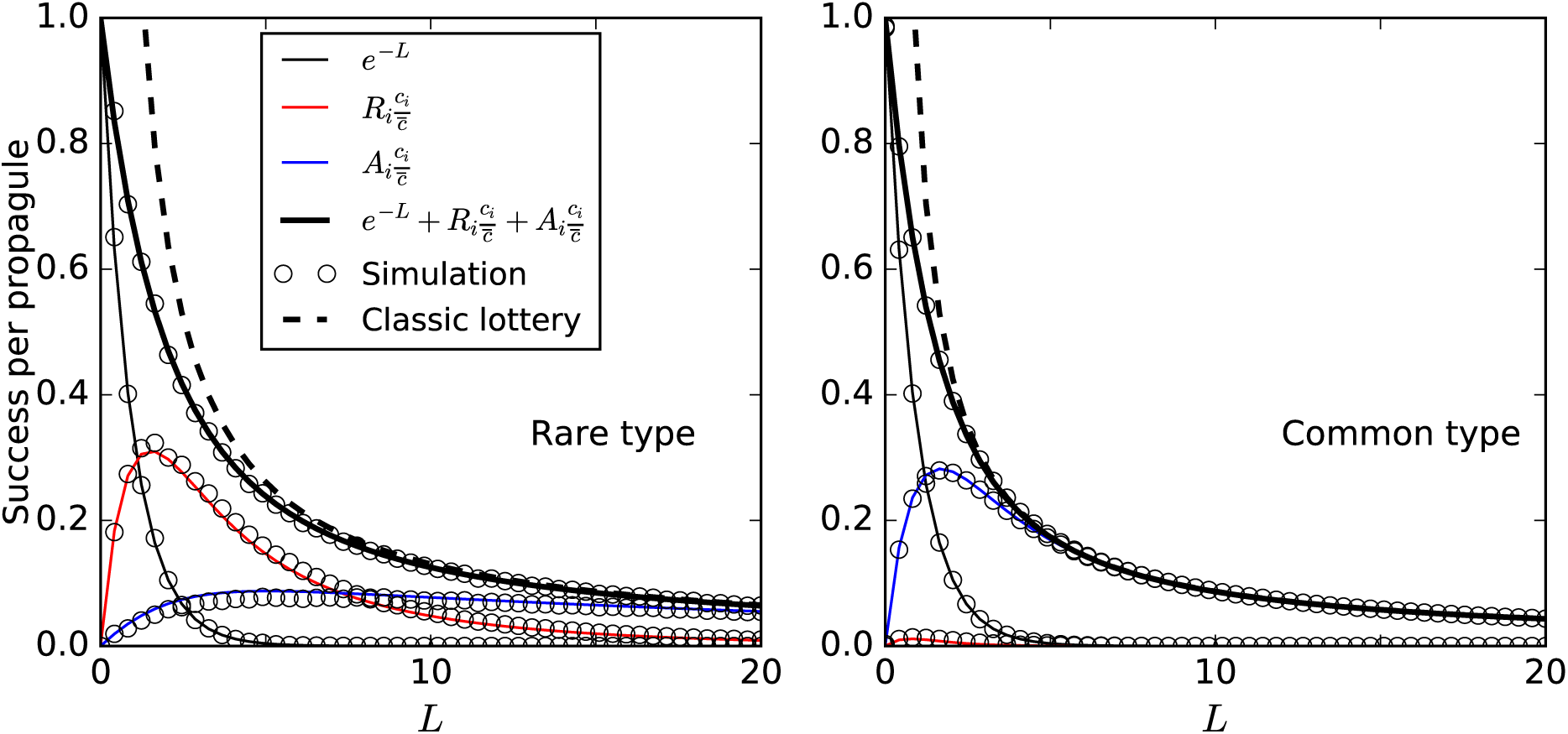
Comparison of Eq. (6), the classic lottery model, and simulations. The vertical axis is per-propagule success rate for all propagules Δ_+_*n*_*i*_*/m*_*i*_, and for the three separate components in Eq. (6). Two types are present with *c*_1_ = 1, *c*_2_ = 1.5 and *l*_2_*/l*_1_ = 0.1. Simulations are conducted as follows: *x*_1_*, x*_2_ values are sampled *U* = 10^5^ times from Poisson distributions with respective means *l*_1_*, l*_2_, and the victorious type in each territory is then decided by random sampling weighted by the lottery win probabilities *c*_*i*_*x*_*i*_*/*(*c*_1_*x*_1_ + *c*_2_*x*_2_). Dashed lines show the failure of the classic lottery model at low density.

In the special case that all types are competitively equivalent (identical *c*_*i*_), Eq. (6) takes a simpler form,

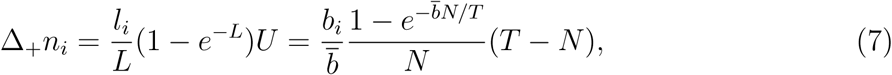

where we have used the fact that 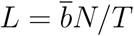 to make the dependence on *b* and *N* explicit (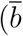 is the population mean *b*). Eq. (7) happens to be exact even though it is a special case of the approximation Eq. (6). This can be deduced directly from Eq. (3): 1 *-e*^*-L*^ is the fraction of territories that receive at least one propagule under Poisson dispersal, (1 *-e*^*-L*^)*U* is the total number of such territories, and type *i* is expected to receive a fraction *l*_*i*_*/L* of these. By similar reasoning, the total number of territories acquired is given by

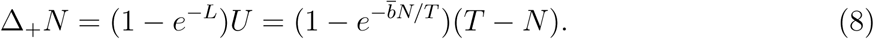

This formula is also exact, but unlike Eq. (7), it also applies when the *c*_*i*_ differ between types.

### Density regulation and selection in the variable-density lottery

Equipped with Eq. (6) we now outline the basic properties of the *b, c* and *d* traits. Adult density *N* is regulated by the birth and mortality rates *b* and *d*; *b* controls the fraction of unoccupied territories that are contested (see Eq. (8)), while *d* controls adult mortality. Competitive ability *c* does not enter Eq. (8), and therefore does not regulate total adult density: *c* only affects the relative likelihood of winning a contested territory.

Selection in our variable-density lottery model is in general density-dependent, by which we mean that the discrete-time selection factor (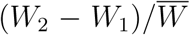) from Eq. (2) may depend on *N.* More specifically, as we show below, *b*- and *c*-selection are density-dependent, but *d*-selection is not. Note that density-dependent selection is sometimes taken to mean a qualitative change in which types are fitter than others at different densities (Travis et al., 2013). While reversal in the order of fitnesses and co-existence driven by density-regulation are possible in our variable-density lottery (a special case of the competition-colonization trade-off; Levins and Culver 1971; Tilman 1994; Bolker and Pacala 1999), questions related to co-existence are tangential to the aims of the current manuscript and will not be pursued further here.

The strength of *b*-selection declines with increasing density. When types differ in *b* only (*b*-selection), Eq. (6) simplifies to Eq. (7), and absolute fitness can be written as 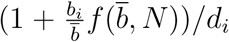 where 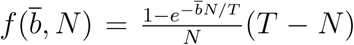 is a decreasing function of *N.* Thus, the selection factor 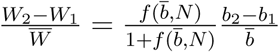 declines with increasing density: the advantage of having greater *b* gets smaller the fewer territories there are to be claimed (Fig. 4).

**Figure 4:**
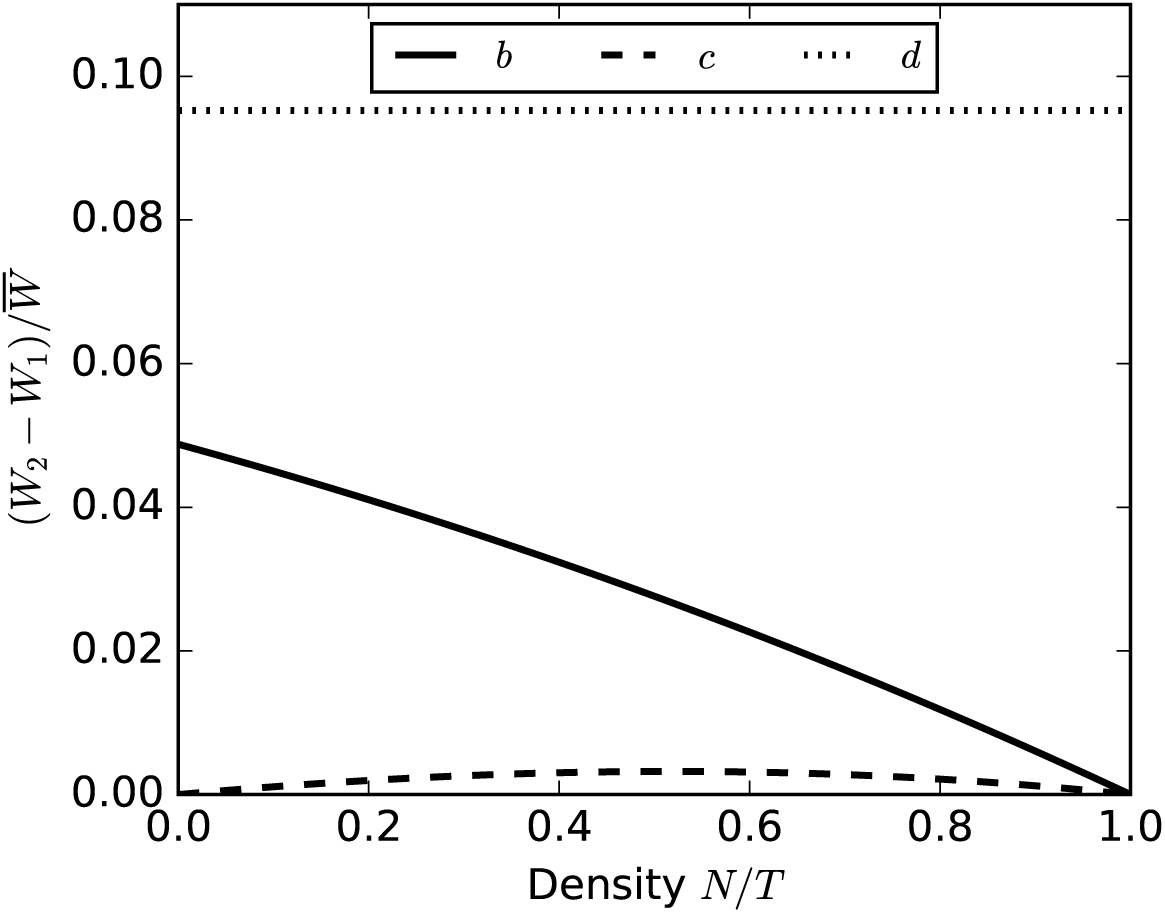
The density-dependence of selection in our variable-density lottery between an adaptive variant 2 and a wildtype variant 1 with at equal frequencies. Here *b*_1_ = 1, *d*_1_ = 2 and *c*_1_ = 1. For *b*-selection we set *b*_2_ = *b*_1_(1+ ϵ), and similarly for *c* and *d*, with ϵ = 0.1. *d*-selection is density-independent, *b*-selection gets weaker with lower territorial availability, while *c*-selection initially increases with density as territorial contests become more important, but eventually also declines as available territories become scarce.

In the case of *c*-selection, Eq. (6) implies that *W*_2_ *-W*_1_ is proportional to 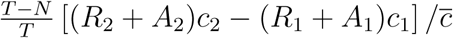 The strength of *c*-selection thus peaks at an intermediate density (Fig. 4), because most territories are claimed without contest at low density (*R*_1_*, R*_2_*, A*_1_*, A*_2_ *→* 0), whereas at high density few unoccupied territories are available to be contested (*T - N →* 0).

Selection on *d* is independent of density, because the density-dependent factor 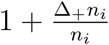
in Eq. (4) is the same for types that differ in *d* only.

### **The response of density to selection;** *c***-selection versus** *K***-selection**

We now turn to the issue of how density changes as a consequence of selection in our variable-density lottery, and in more familiar models of selection in density-regulated populations. In the latter, selection under crowded conditions typically induces changes in equilibrium density (see Introduction). In our variable-density lottery model, however, the competitive ability trait *c* is not density-regulating, even though *c* contributes to fitness under crowded conditions. Consequently, *c*-selection does not cause density to change. In this section we compare this *c*-selection behavior with the previous literature, which we take to be exemplified by MacArthur’s *K*-selection argument (MacArthur and Wilson, 1967).

MacArthur considered two types (with densities *n*_1_ and *n*_2_) in a constant environment subject to density-dependent growth,

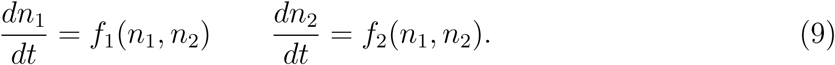

The outcome of selection is determined by the relationship between the nullclines *f*_1_(*n*_1_*, n*_2_) = 0 and *f*_2_(*n*_1_*, n*_2_) = 0. Specifically, a type will be excluded if its nullcline is completely contained in the region bounded by the other type’s nullcline.

MacArthur used the four intersection points of the nullclines with the axes, defined by *f*_1_(*K*_11_, 0) = 0, *f*_1_(0*, K*_12_) = 0, *f*_2_(*K*_21_, 0) = 0 and *f*_2_(0*, K*_22_) = 0, to analyze each type’s exclusion or persistence. Note that only *K*_11_ and *K*_22_ are equilibrium densities akin to the *K* parameter in the logistic model; the other intersection points, *K*_12_ and *K*_21_, are related to competition between types. For instance, in the Lotka-Volterra competition model we have

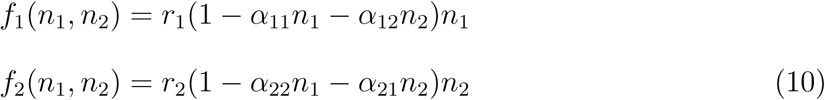

where *α*_11_ = 1*/K*_11_ and *α*_22_ = 1*/K*_22_ measure competitive effects within types, while *α*_12_ = 1*/K*_12_ and *α*_21_ = 1*/K*_21_ measure competitive effects between types. Hence, “fitness is *K*” in crowded populations (MacArthur and Wilson, 1967, pp. 149) in the sense that selection either favors the ability to keep growing at ever higher densities (moving a type’s own nullcline outwards), or the ability to suppress the growth of competitors at lower densities (moving the nullcline of competitors inwards; Gill 1974). However, even if the initial and final densities of an adaptive sweep in the Lotka-Volterra model are the same, *N* nevertheless does change transiently (Fig. 5a). Constant-*N* over a sweep only occurs for a highly restricted subset of *r* and *α* values (Appendix B; Mallet 2012; Gill 1974; Smouse 1976).

**Figure 5:**
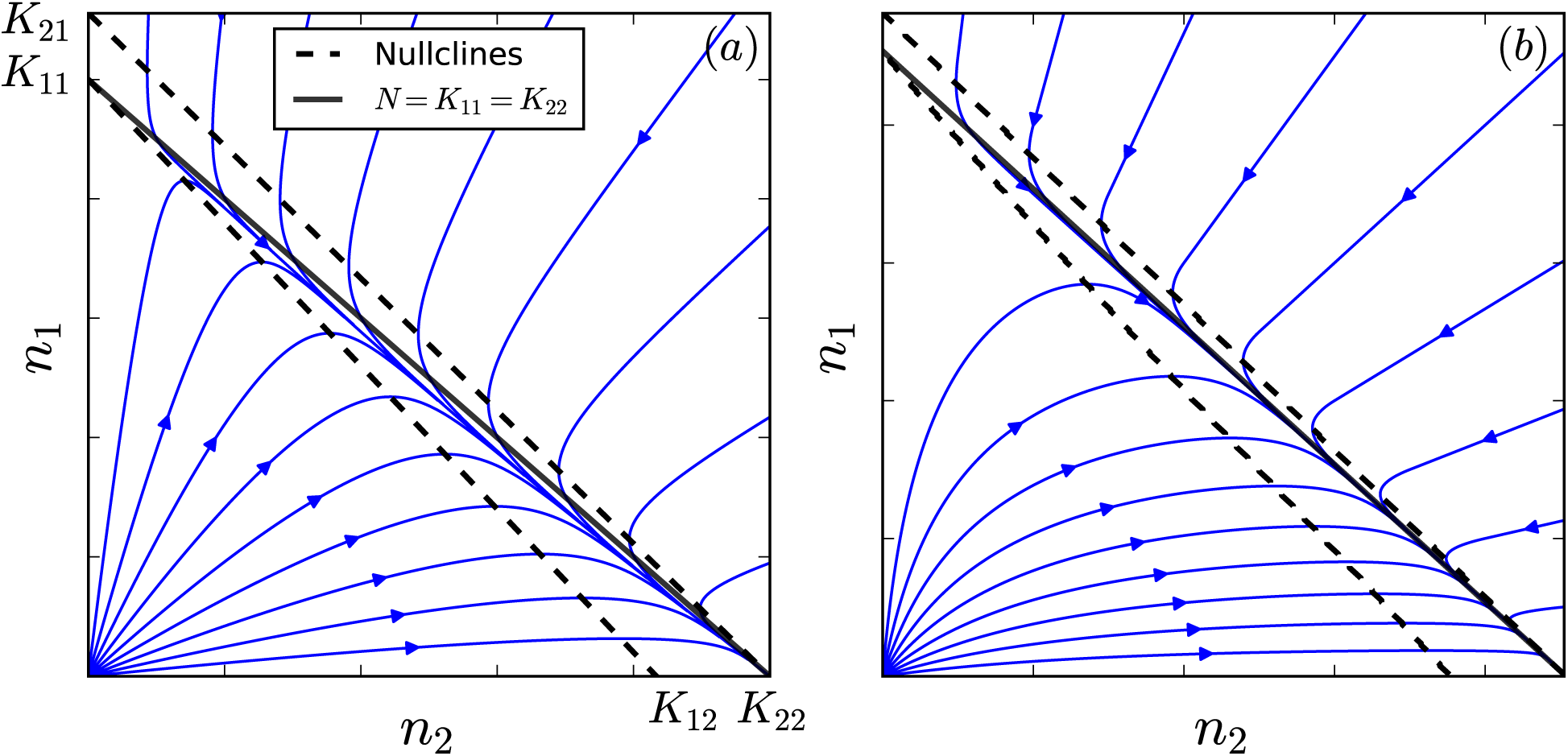
Selection between types with identical equilibrium density but different inter-type competitive ability. (a) Lotka-Volterra competition (Eq. 10) with *r*_1_ = *r*_2_ = 1, *α*_11_ = *α*_22_ = 1, *α*_12_ = 0.9 and *α*_21_ = 1.2. Trajectories do not follow the line *N* = *K*_11_ = *K*_22_. (b) Lottery competition (Eq. 6) with *b*_1_ = *b*_2_ = 5, *d*_1_ = *d*_2_ = 1.1 and *c*_1_*/c*_2_ = 5. The lottery model nullclines are defined by *W*_1_ = 1 (lower nullcline) and *W*_2_ = 1 respectively in Eq. (4). For the discrete-time Eq. (6), trajectories are the flow lines of the vector field obtained by evaluating the direction of the local changes in *n*_1_ and *n*_2_; these converge on the line *N* = *K*_11_ = *K*_22_.

In contrast, density trajectories for *c*-selection in our variable-density lottery converge on a line of constant equilibrium density (Fig. 5b). This means that once *N* reaches demographic equilibrium, selective sweeps behave indistinguishably from a constant-*N* relative fitness model(Fig. 1b). Thus, for *c*-sweeps in a constant environment, the selection factor (*W*_2_ 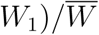 in Eq. (2) is density-independent. This uncoupling of density from *c*-selection arises due to the presence of an excess of propagules which pay the cost of selection without affecting adult density (Nei, 1971).

### Density-regulating traits under strong selection

For density to matter in Eq. (2), selection must be density-dependent and density must be changing. This can occur in a constant environment if selection acts on a density-regulating trait. Consider the simple birth-death model (Kostitzin, 1939)

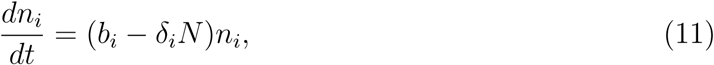

where *δ*_*i*_ is per-capita mortality due to crowding. Starting from a type 1 population in equilibrium, a variant with *δ*_2_ = *δ*_1_(1 −ϵ) has density-dependent selection coefficient *s* = ϵ *δ* _1_*N*, which will change over the course of the sweep as *N* shifts from its initial type 1 equilibrium to a type 2 equilibrium. The equilibrium densities at the beginning and end of the sweep are *N*_initial_ = *b*_1_*/δ*_1_ and *N*_final_ = *b*_1_*/*(*δ*_1_(1 −ϵ)) = *N*_initial_*/*(1 −ϵ) respectively, and so *s*_initial_ = ϵ *b*_1_ and *s*_final_ = *s*_initial_*/*(1 −ϵ). Consequently, substantial deviations from Eq. (1) occur if there is sufficiently strong selection on *δ* (Fig. 6; Kimura 1978; Kimura and Crow 1969).

**Figure 6:**
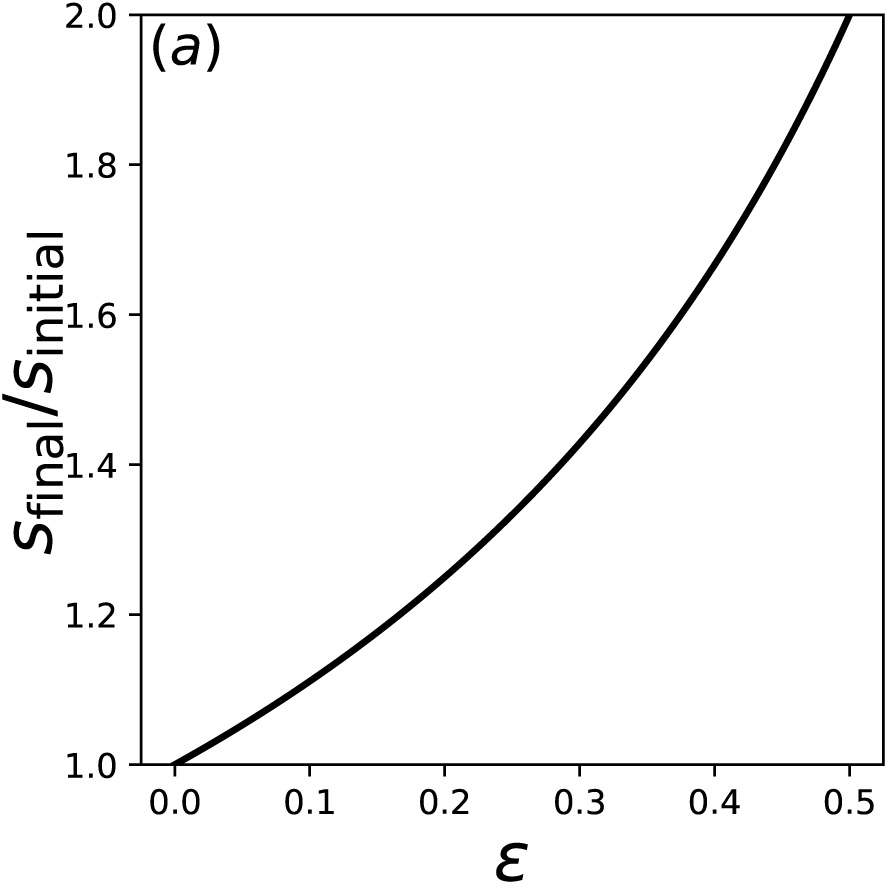
Change in the selection coefficient between the beginning and end of a sweep of a type that experiences proportionally 1 − ϵ fewer crowding-induced deaths. The population is in demographic equilibrium at the start and end of the sweep.

In our variable density lottery, *b* regulates density and is subject to density-dependent selection, yet *b*-sweeps are qualitatively different from *δ* sweeps in the above example. Greater *b* means not only that more propagules contest the available territories, but also that a greater fraction of unoccupied territories receive propagules. Together, the net density-dependent effect on *b*-selection is negligible: in a single-type equilibrium we have *W*_*i*_ = 1 and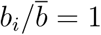, and hence the density-dependence factor 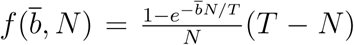 in Eq. (7) has the same value *d*_*i*_ − 1 at the beginning and end of a *b*-sweep (recall that 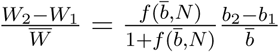 for *b*-selection). During the sweep there is some deviation in 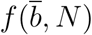, but this deviation is an order of magnitude smaller than for a *δ*-sweep (the density-dependent deviation in Fig. 6 is of order ϵ, whereas the analogous effect for a *b*-sweep in our variable-density lottery is only of order ϵ ^2^; see Appendix C for details). Since selection must already be strong for a *δ*-sweep to invalidate Eq. (1), the density-independent model applies almost exactly for equilibrium *b*-sweeps (Fig. 7).

**Figure 7:**
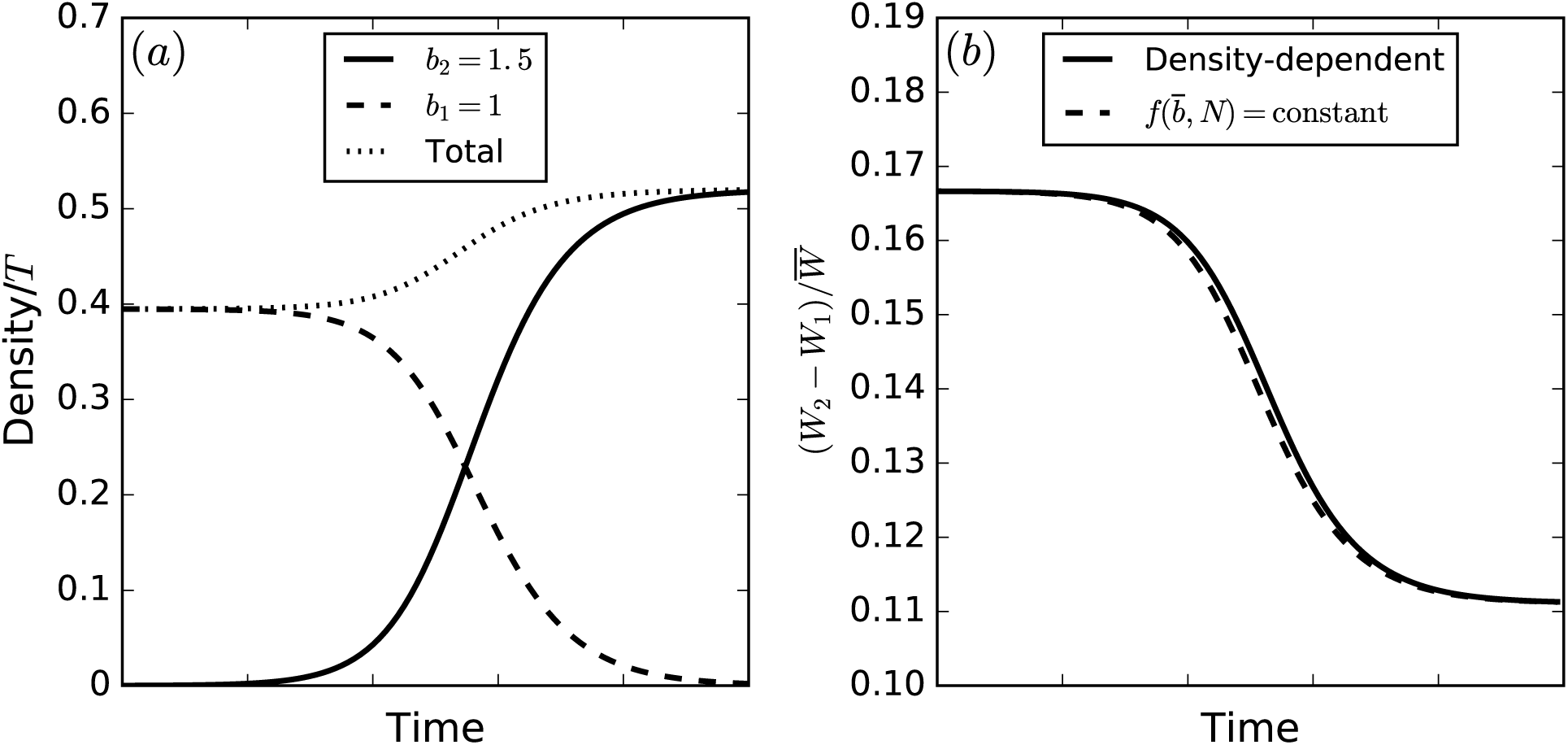
Equilibrium *b*-sweeps behave as though selection is independent of density even though *b*-selection is density-dependent in general. Panel (b) shows the density-dependent selection factor 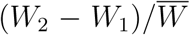 predicted by E<u>q</u>. (6) (solid line) compared to the same selection factor with the density-dependence term 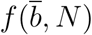 held constant at its initial value (dashed line).

However, if selection acts simultaneously on more than one trait in our variable-density lottery, then evolution in a density-regulating trait can drive changes in the strength of selection on another trait subject to density-dependent selection. For instance, if selection acts simultaneously on *b* and *d*, then 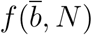 changes value from *d*_1_ − 1 to *d*_2_ − 1 over a sweep. The dynamics of density will then affect the selection factor 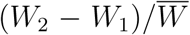 and cause deviations analogous to selection on *δ* in the continuous time case (Fig. 8).

**Figure 8:**
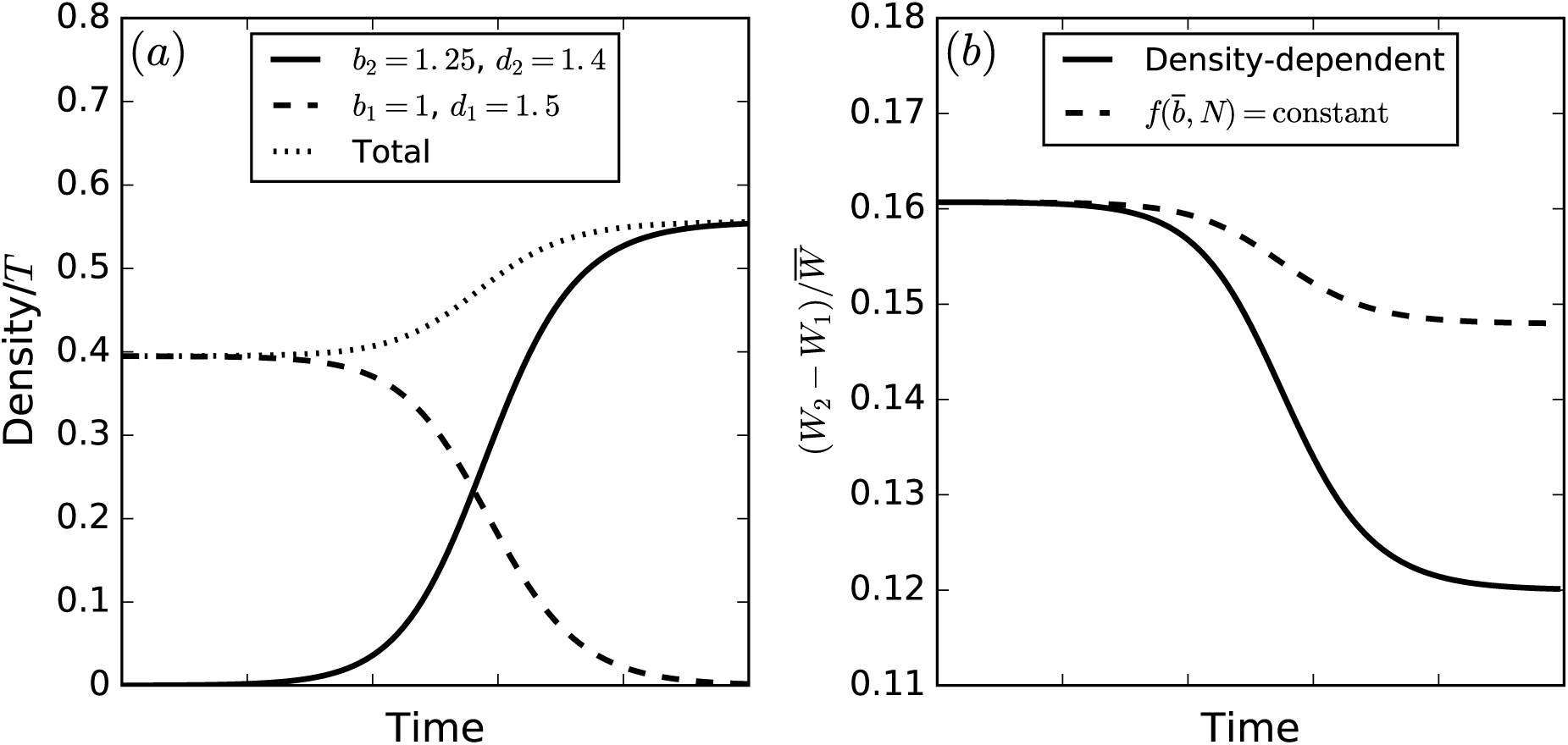
Simultaneous selection on *b* and *d* induces density-dependence in the selection factor 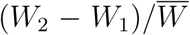 Panel (b) shows the predictions of Eq. (7) (solid line) versus the same with the density-dependence factor 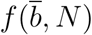 held constant at its initial value.

## Discussion

Summarizing the properties of selection in our variable-density lottery model: (i) *c*-selection is density-dependent, but *c* does not regulate density; (ii) *d* regulates density, but *d*-selection is density-independent; (iii) *b* regulates density and *b*-selection is density-dependent. Yet, despite the differences between *b, c* and *d*, selection in a constant environment that only involves one of these traits obeys the density-independent relative fitness description of selection almost exactly (that is, 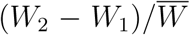 in Eq. (2) is approximately independent of density). This density-independence breaks down when strong selection acts on more than one of *b, c* and *d* (Fig. 8). The *c* and *d* traits exemplify the two distinct directions in which density and selection can interact: selection may depend on density, and density may change in response to ongoing selection (Prout, 1980). The combination of both is necessary to invalidate the constant-*s* approximation. Remarkably, the *b* trait demonstrates that the combination is not sufficient; the density-dependence of *b*-selection effectively disappears over equilibrium-to-equilibrium *b*-sweeps.

The distinctive properties of selection in the variable-density lottery arise from a reproductive excess which appears when the number of propagules is greater than the number of available territories. Then only *≈* 1*/L* of the juveniles contesting unoccupied territories survive to adulthood. Unlike the role of adult density *n*_*i*_ in single-life-stage models, it is the propagule densities *l*_*i*_ that represent the crowding that drives competition. Reproductive excess produces relative contests in which fitter types grow at the expense of others by preferentially filling the available adult “slots”. The number of available slots can remain fixed or change independently of selection at the juvenile stage. By ignoring reproductive excess, single life-stage models are biased to have total population density be more sensitive to ongoing selection. In this respect, the viability selection heuristics that are common in population genetics (Gillespie, 2010, pp. 61) actually capture an important ecological process without making the full leap to complex age-structured models.

Traits like *b, c* and *d* will often have pleiotropic interactions which mean that adaptive sweeps will behave similarly to the familiar “*δ* sweeps” (”Density regulating traits under strong selection”). Thus, our analysis of the variable density lottery does not necessarily imply that Eq. (1) and (2) should apply more broadly than previously thought. Rather, *b, c* and *d* represent a possible set of idealized fitness components mediating the interplay between selection and density in density-regulated populations. The conceptual distinctions they highlight will hopefully be useful regardless of the biological realism of our variable density lottery model.

Apart from familiar examples such as reef fishes (Chesson and Warner, 1981), many biological systems do not obviously satisfy the assumptions of our variable density lottery. However, even if competition occurs primarily via consumable resource exploitation, spatial localization of consumable resources (e.g. for plants due to restricted movement of nutrients through soils) will tend to create territorial contests similar to the lottery model, where resource competition only occurs locally and can be sensitive to contingencies such as the timing of propagule arrival (Bolker and Pacala, 1999). In this case, resource competition is effectively subsumed into a territorial competitive ability trait akin to *c*, which would likely affect *N* much more weakly than suggested by the *R\** rule (assuming no pleiotropic interactions with *b* or *d*). Moreover, even in well-mixed populations, competition does not only involve indirect exploitation of shared resources, but also direct interference. Interference competition can dramatically alter the dynamics of resource exploitation (Case and Gilpin, 1974; Amarasekare, 2002), and is more likely than the exploitation of shared resource pools to involve relative contests akin to *c*-selection. For instance, sexual selection can be viewed as a form of relative interference competition between genotypes.

In the analysis presented here we have restricted our attention to selection in demographically stable populations. The largest deviations from the approximation of density-independent selection (as represented by Eqs. (1) and (2)) will likely occur in populations far from demographic equilibrium e.g. as a result of a temporally-variable environment. This is because extremely strong selection is needed to change population density by an amount comparable to environmental varability (see Fig. 6). By contrast, temporally-variable environments can dramatically alter frequency trajectories for individual sweeps (e.g. Fig. 9.5 in Otto and Day (2011); Fig. 5 in Mallet (2012)), as well as the long-term outcomes of selection (Lande et al., 2009).

**Figure 9:**
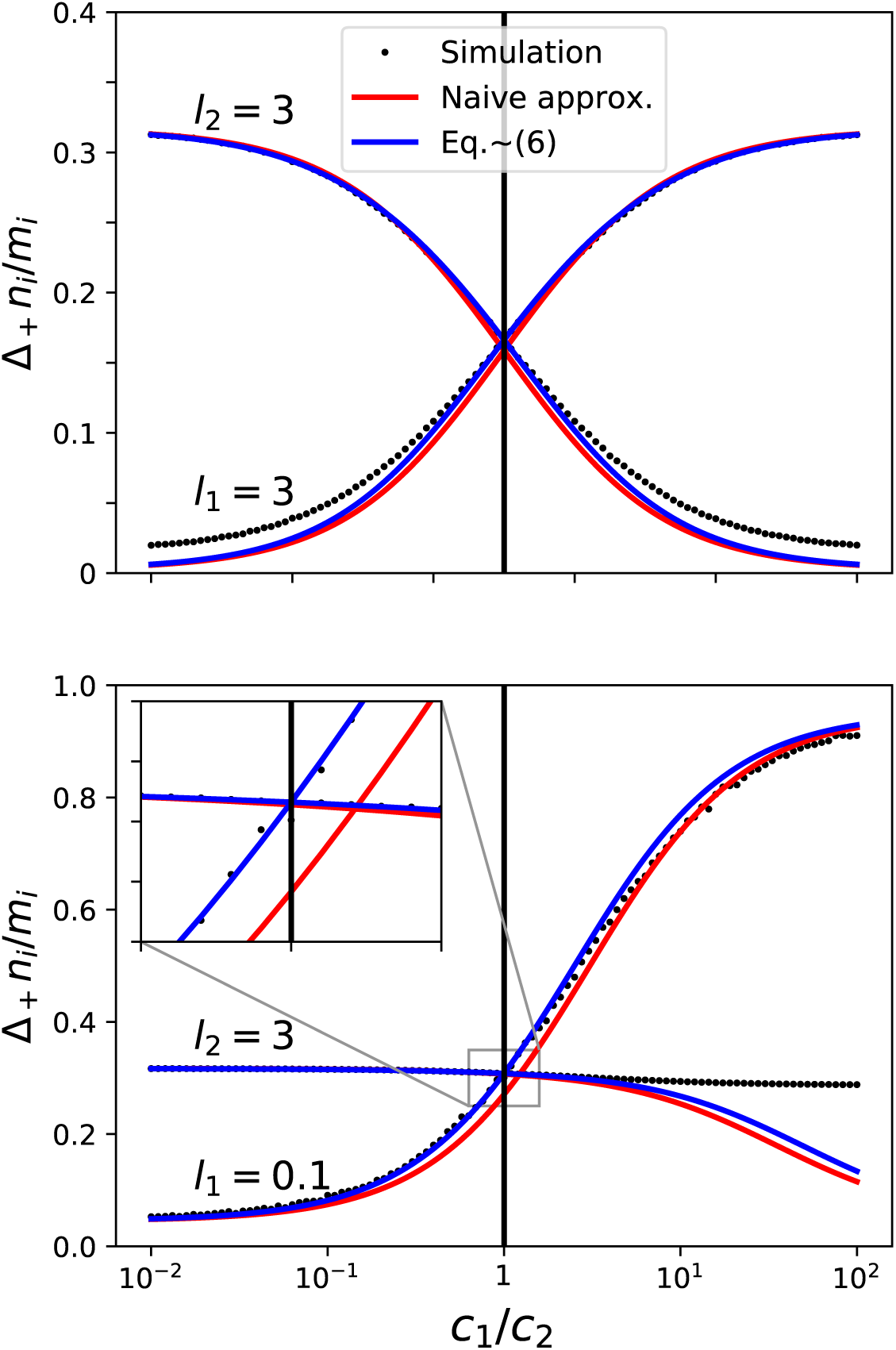
Comparison of our effective mean approximation Eq. (6) with simulations, and also with the naive ⟨ ⟩_*r*_ = ⟨ ⟩_*p*(**x***|x*_*i*_=1*,X*_*i*_*≥*1)_ and ⟨ ⟩ _*a*_ = ⟨ ⟩ _*p*(**x***|x*_*i*_*≥*2)_ approximation, as a function of the relative *c* difference between two types. Eq. (6) breaks down in the presence of large *c* differences. The inset shows the pathology of the naive approximation — growth rates for rare and common types are not equal in the neutral case *c*_1_ = *c*_2_. Simulation procedure is the same as in Fig. 3, with *U* = 10^5^.

This suggests that in systems like the wild *Drosophila* example mentioned in the Introduction, there may indeed be no choice but to abandon relative fitness. Our variable-density lottery could provide a useful starting point for analyzing evolution in this and other farfrom-equilibrium situations for two reasons: 1) the *b, c, d* trait scheme neatly distinguishes between different aspects of the interplay between density and selection; 2) lottery models in general are mathematically similar to the Wright-Fisher model, which should facilitate the analysis of genetic drift when *N* is unstable.

## Acknowledgments

We thank Peter Chesson and Joachim Hermisson for many constructive comments on an earlier and quite different version of this manuscript. This work was financially supported by the National Science Foundation (DEB-1348262) and the John Templeton Foundation (60814).

## Appendix A: Growth equation derivation

Here we derive Eq. (6). Following the notation in the main text, the Poisson distributions for the *x*_*i*_ (or some subset of the *x*_*i*_) will be denoted *p*, and we use *P* as a general shorthand for the probability of particular outcomes. We denote the vector of propagule abundances by **x** = (*x*_1_*,…, x*_*G*_) in a given territory, and the analogous vector of nonfocal abundances by **x**_*i*_ = (*x*_1_*,…, x*_*i−*1_*, x*_*i*+1_… *, x*_*G*_). The corresponding total propagule numbers are denoted *X* = Σ _*j*_ *x*_*j*_ and *X*_*i*_ = *X - x*_*i*_.

Similar to the classic lottery model, our approximation involves replacing the *x*_*i*_ with effective mean values. However, as discussed in the text preceding Eq. (6), it is important to treat the *x*_*i*_ = 1 case separately when allowing for low propagule densities. We thus start by separating the right hand side of Eq. (3) into three components

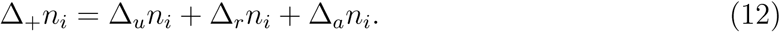

The relative magnitude of these components depends on the propagule densities *l*_*i*_. The first component, Δ_*u*_*n*_*i*_, accounts for territories where only one focal propagule is present (*x*_*i*_ = 1 and *x*_*j*_ = 0 for *j ≠ i*; *u* stands for “uncontested”). The proportion of territories where this occurs is *l*_*i*_*e*^*-L*^, and so

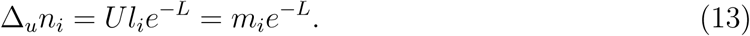

The second component, Δ_*r*_*n*_*i*_, accounts for territories where a single focal propagule is present along with at least one non-focal propagule (*r* stands for “rare”). The number of territories where this occurs is *Up*_*i*_(1)*P* (*X*_*i*_ *≥* 1) = *m*_*i*_*e*^*-l*^_*i*_ (1 *- e*^−(*L-l*^_*i*_)). Thus

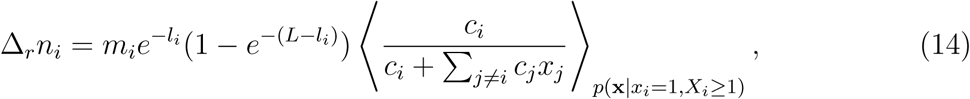

where ⟨ ⟩ _*p*(**x***|x*_*i*_=1*,X*_*i*_*≥*1)_ denotes the expectation with respect to the conditional probability distribution *p*(**x***|x*_*i*_ = 1*, X*_*i*_ *≥* 1) of propagule abundances in those territories where exactly one focal propagule, and at least one non-focal propagule, landed.

The final contribution, Δ_*a*_*n*_*i*_, accounts for territories where two or more focal propagules are present (*a* stands for “abundant”). Similar to Eq. (14), we have

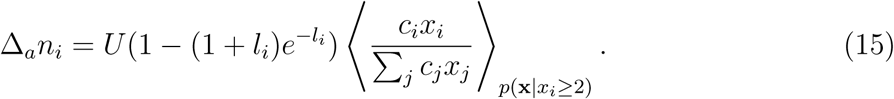

To derive Eq. (6) we approximate the expectations in Eq. (14) and Eq. (15) by replacing *x*_*i*_ and the *x*_*j*_ with “effective” mean values as follows

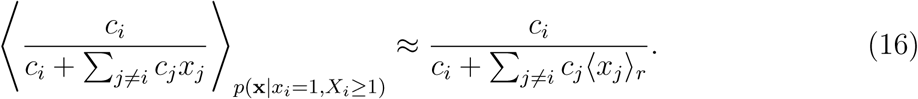

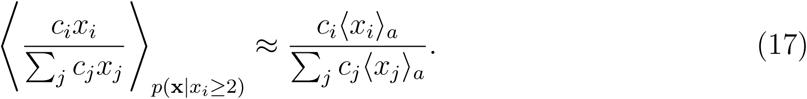

Here ⟨ ⟩_*r*_ and ⟨ ⟩ _*a*_ are the effective means, which are defined in the following subsection.

### **The effective means** ⟨ ⟩_*r*_ **and** ⟨ ⟩_*a*_

The decomposition Eq. (12) is exact and involves no additional assumptions. However this decomposition complicates our approximation procedure because the separate components in Eq. (12) must be approximated in a consistent manner.

To illustrate this consistency requirement, suppose that two identical types (same *b, c* and *d*) are present, the first with small density *l*_1_ ⟪ 1 and the second with large density *l*_2_ ⟫ 1. In this case, uncontested territories make up a negligible fraction of *U*; the first type’s territorial acquisition is almost entirely due to Δ_*r*_*n*_1_; and the second type’s territorial acquisition is almost entirely due to Δ_*a*_*n*_2_. For consistency, the approximate per-capita growth rates in (16) and (17) must be equal Δ_*r*_*n*_1_*/m*_1_ = Δ_*a*_*n*_2_*/m*_2_. Even small violations of this consistency condition would mean exponential growth of one type relative to the other. This behavior is pathological, because any single-type population can be arbitrarily partitioned into identical rare and common subtypes. Thus, predicted growth or decline would depend on an arbitrary assignment of rarity.

Suppose that we naively used the conditional distributions *p*(**x***|x*_*i*_ = 1*, X*_*i*_ *≥* 1) and *p*(**x***|x*_*i*_ *≥* 2) to calculate the effective means, such that ⟨ ⟩ _*r*_ = ⟨ ⟩ _*p*(**x***|x*_*i*_=1*,X*_*i*_*≥*1)_ and ⟨ ⟩ _*a*_ = ⟨ ⟩ _*p*(**x***|x*_*i*_*≥*2)_. Then, in the example from the previous paragraph (*l*_1_ ⟪ 1, *l*_2_ ⟫ 1), the right hand side of Eq. (16) would be *≈* 1*/*(*l*_2_ + 1), and so Δ_*r*_*n*_1_*/m*_1_ *≈* 1*/*(*l*_2_ + 1) in Eq. (14). Similarly, Σ _*j*_⟨ *x*_*j*_ ⟩ _*a*_ *≈ l*_2_ in Eq. (17), and so Δ_*a*_*n*_2_*/m*_2_ *≈* 1*/l*_2_. Thus, the rare type would be predicted to decline in frequency even though it has identical traits.

This pathological behavior occurs because the expected total density of propagules in the respective groups of territories are different: ⟨ *X* ⟩ _*p*(**x***|x*_1_=1*,X*_1_*≥*1)_ *≈ l*_2_ + 1 *>* ⟨ *X*⟩ _*p*(**x***|x*_2_*≥*2)_ *≈ l*_2_. As a result, the rare type’s behavior is approximated as though it experiences more intense lottery competition than the common type, which cannot be the case since the two types are identical. The effective means must thus be taken in a way that ensures that the expected total propagule density is the same in Eq. (16) and Eq. (17).

We achieve this as follows. For nonfocal types *j ≠ i*, we separately evaluate the *X*-dependence of the conditional dispersal probabilities to ensure that *X* has the same distribution for both ⟨ *X*⟩ _*r*_ and ⟨ *X*⟩ _*a*_. Specifically, we assume that *X* follows a Poisson distribution with rate parameter *L*, conditional on *X ≥* 2; this distribution will be denoted *P* (*X|X ≥* 2). However, for the focal type *i*, we use the exact conditional dispersal distributions *p* to calculate the effective mean,

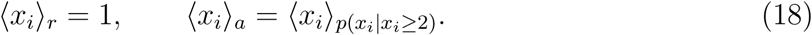

As we will see, these effective means are straightforward to calculate analytically, and ensure that the expected total propagule density ⟨ *x*_*i*_ *X*⟩ + Σ _*j/*=*i*_ ⟨*x*_*j*_⟩ is the same in Eq. (16) and Eq. (17).

Starting with Eq. (16), we only need to evaluate ⟨*x*_*j*_⟩ _*r*_ since ⟨*x*_*i*_⟩ _*r*_ = 1. To evaluate the *X*-dependence separately, we first hold *X* fixed to obtain

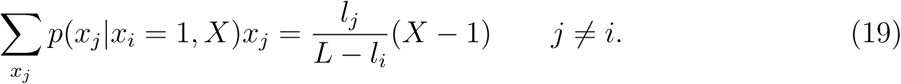

The right hand side is obtained by observing that the sum on the left is the expected number of propagules with type *j* that will be found in a territory which received *X −* 1 nonfocal propagules in total. We then take the expectation with respect to *P* (*X|X ≥* 2) to give

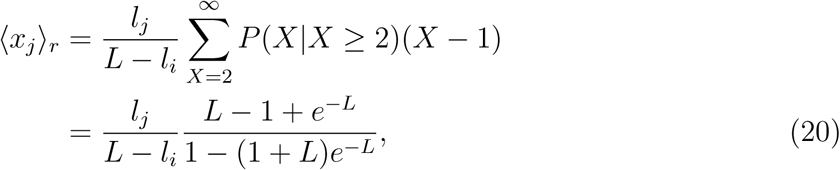

where the last line follows from 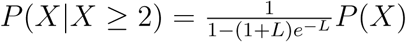 and 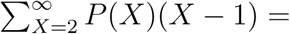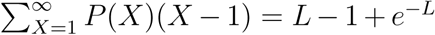. Substituting Eqs. (16) and (20) into Eq. (14), we obtain

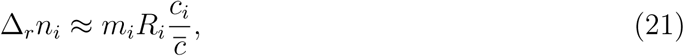

where *R*_*i*_ is defined in Eq. (7).

Turning now to Eq. (17), from Eq. (18) the mean focal abundance is

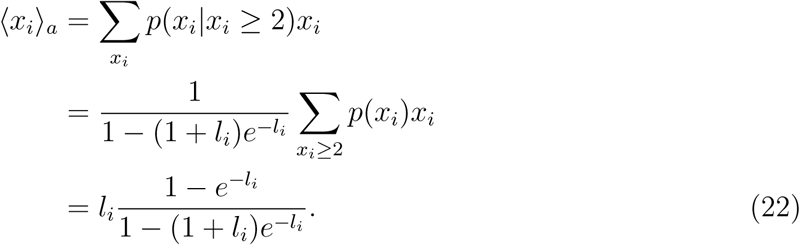

For nonfocal types *j ≠ i*, we have analogously to Eq. (19),

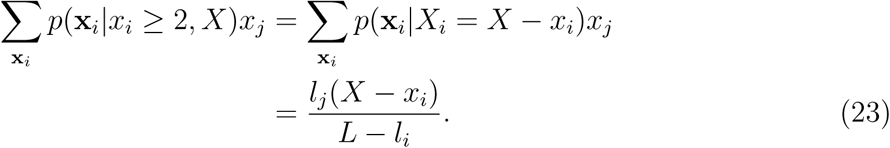

Again taking the expectation with respect to *P* (*X|X ≥* 2) yields

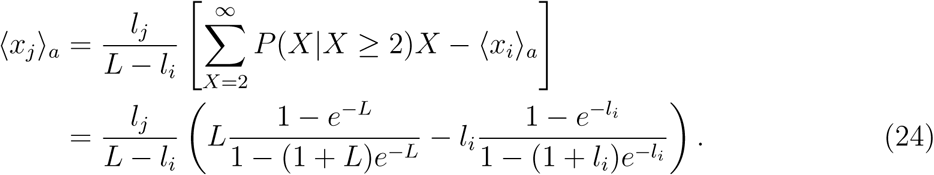

Combining these results with Eqs. (15) and (17), we obtain

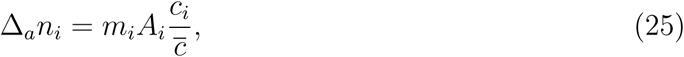

where *A*_*i*_ is defined in Eq. (7).

It is easily verified from Eqs. (20), (22) and (24) that the total expected propagule density is the same in in Eq. (16) and Eq. (17) i.e. ⟨ *x*_*i*_ ⟩ _*r*_ + Σ_*j i*_ *x*_*j*_⟨ ⟩ _*r*_ = *x*_*i*_⟨ ⟩ _*a*_ + *x*_*j*_⟨ ⟩_*a*_ = ⟨*X*⟩_*P* (*X|X≥*2)_. As a result, Eq. (6) satisfies the consistency requirement (see Fig. 9).

### Approximation limits

Having derived the approximation Eq. (6), we now evaluate its domain of validity. Eq. (6) relies on ignoring the fluctuations in *x*_*i*_ and *x*_*j*_, such that we can replace them with constant effective mean values. To justify this, we show that the standard deviations *σ*_*p*(**x***|x*_*i*_=1*,X*_*i*_*≥*1)_ (Σ _*j≠i*_ *c*_*j*_*x*_*j*_) and *σ* **_*p*(**x***|x*__*i*_*≥*2)**(Σ _*j*_ *c*_*j*_*x*_*j*_) are small compared to the corresponding means (Σ _*j≠i*_ *c*_*j*_*x*_*j*_) _*p*(**x***|x*_*i*_=1*,X*_*i*_*≥*1)_ and (Σ _*j*_ *c*_*j*_*x*_*j*_) **_*p*(**x***|x*__*i*_*≥*2)** in Eqs. (16) and (17). This result means that using the exact distributions *p*(**x***|x*_*i*_ = 1*, X*_*i*_ *≥* 1) and *p*(**x***|x*_*i*_ *≥* 2) for the effective means would produce an accurate approximation of the components in (12) (though, as we have seen, not a consistent one). It is then clear that the effective means derived in the previous section will also give an accurate approximation since their magnitudes are similar to the exact means; this is obvious from the fact that the expected total number of propagules is of order max{*L,* 2} in both cases.

We first consider the means and standard deviations in Eq. (16). We have *⟨x*_*j*_*⟩* _*p*(**x***|x*_*i*_=1*,X*_*i*_*≥*1)_ = *l*_*j*_*/C*, where *C* = 1 *- e*^−(*L-l*_*i*_)^, and the corresponding variances and covariances are given by

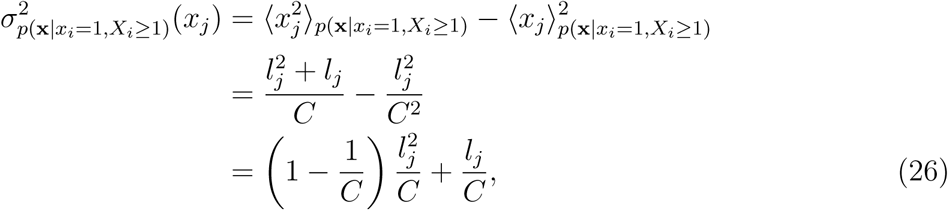

and

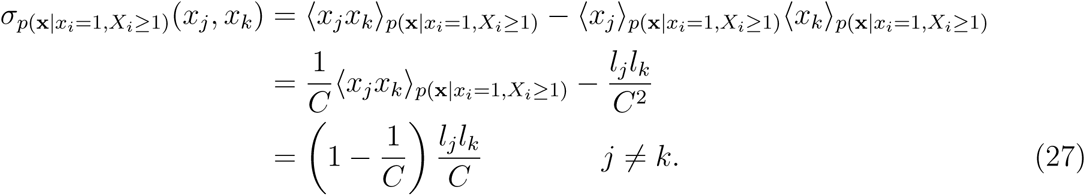

Note that 1 − 1*/C* is negative because *C <* 1. Decomposing the variance in Σ_*j≠i*_ *c*_*j*_*x*_*j*_,

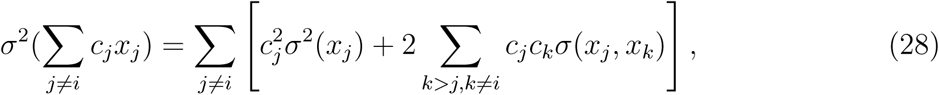

we obtain

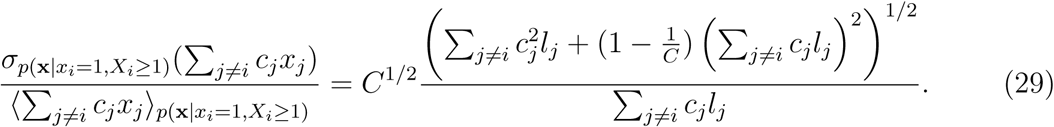

Eq. (29) shows that, when the *c*_*j*_ have similar magnitudes (their ratios are of order one), Eq. (16) is an excellent approximation. The right hand side of Eq. (29) is then approximately equal to 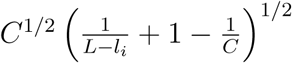, which is small for both low and high nonfocal densities. The worst case scenario occurs when *L - l*_*i*_ is of order one, and it can be directly verified that Eq. (16) is then still a good approximation (see Fig. 9).

Turning to Eq. (17), all covariances between nonfocal types are now zero, so that.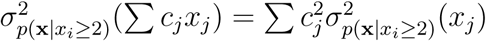 For nonfocal types 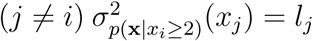 whereas for the focal type we have

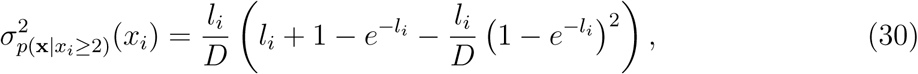

where *D* = 1 − (1 + *l*_*i*_)*e*^*-l*^^*i*^, and

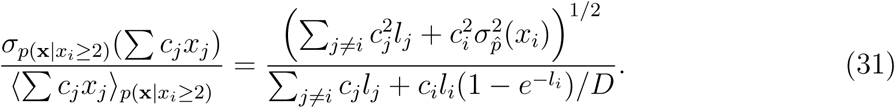

Similarly to Eq. (29), the right hand side of Eq. (31) is small for both low and high nonfocal densities provided that the *c*_*j*_ have similar magnitudes. Again, the worst case scenario occurs when *l*_*i*_ and *L - l*_*i*_ are of order 1, but Eq. (17) is still a good approximation in this case (Fig. 9).

In both Eqs. (29) and (31), the standard deviation in Σ_*≠i*_ *c*_*j*_*x*_*j*_ can be large relative to its mean if some of the *c*_*j*_ are much larger than the others. Specifically, in the presence of a rare, strong competitor (*c*_*j*_*l*_*j*_ *≫ c*_*j*_ *l*_*j*_ for all other nonfocal types *j*^*t*^, and *l*_*j*_ *≪* 1), then the right hand side of Eqs. (29) and (31) can be large and we cannot make the replacement Eq. (16). Fig. 9 shows the breakdown of the effective mean approximation when there are large differences in *c*.

## Appendix B: Total density under Lotka-Volterra competition

Here we show that under the Lotka-Volterra model of competition, total density *N* does not in general remain constant over a selective sweep in a crowded population even if the types have the same equilibrium density (for a related discussion on the density- and frequency-dependence of selection in the Lotka-Volterra model, see (Smouse, 1976; Mallet, 2012)).

We assume equal effects of crowding within types *α*_11_ = *α*_22_ = *α*_intra_ and *N* = 1*/α*_intra_ and check whether it is then possible for 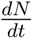 to be zero in the sweep (*n*_1_*, n*_2_ ≠ 0). Substituting these conditions into Eq. (10), we obtain

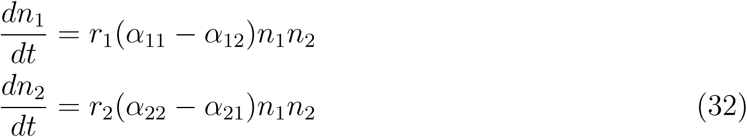

Adding these together, 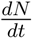 can only be zero if

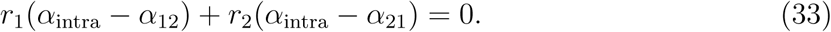

To get some intuition for Eq. (33), suppose that a mutant arises with improved competitive ability but identical intrinsic growth rate and equilibrium density (*r*_1_ = *r*_2_ and *α* _11_ = *α* _22_). This could represent a mutation to an interference competition trait, for example (Gill, 1974). Then, according the above condition, for *N* to remain constant over the sweep, the mutant must find the wildtype more tolerable than itself by exactly the same amount that the wildtype finds the mutant less tolerable than itself.

Even if we persuaded ourselves that this balance of inter-type interactions is plausible in some circumstances, when multiple types are present the requirement for constant *N* becomes

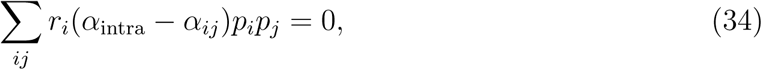

which depends on frequency and thus cannot be satisfied in general for constant inter-type coefficients *α*_*ij*_. Therefore, Lotka-Volterra selection will generally involve non-constant *N.*

## **Appendix C: Density-dependence of** *b***-selection**

In section “Density-regulating traits under strong selection” we argued that the density-dependent factor 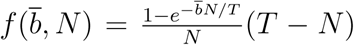 is unchanged at the beginning and end points of an equilibrium to equilibrium sweep of a type with higher *b*. Here we estimate the magnitude of the deviation in 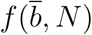 during the sweep.

For simplicity, we introduce the notation *D* = *N/T* and assume that *D* is small. We can thus make the approximation 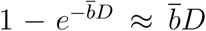 and 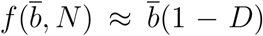 We expect this to be a conservative approximation based on the worst case scenario, because *N* is most sensitive to an increase in *b* in this low-density linear regime. We first calculate the value of 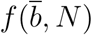 at the halfway point in a sweep, where the halfway point is estimated with simple linear averages for *b* and *N.* The sweep is driven by a *b* variant with *b*_2_ = *b*_1_(1 + ϵ), and we denote the initial and final densities by *D*_1_ and *D*_2_ respectively, where we have *f*_initial_ = *b*_1_(1 *- D*_1_) = *d*_1_ − 1 = *f*_final_ = *b*_2_(1 *- D*_2_). We obtain

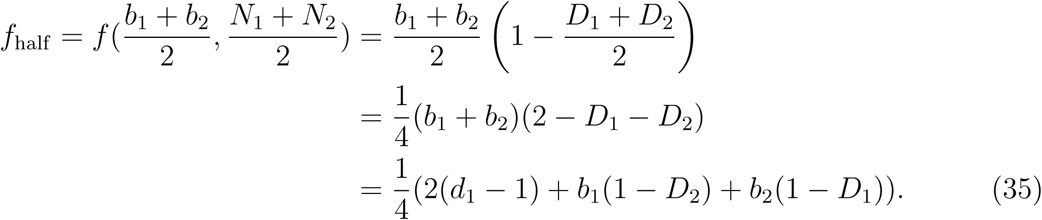

Dividing by *d*_1_ − 1, the proportional deviation in *f* (*N)* at the midpoint of the sweep is

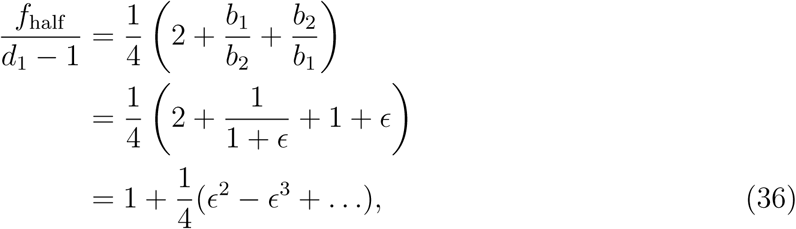

where we have used the Taylor expansion 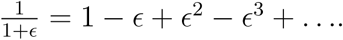

By contrast, for a *δ* sweep in Eq. (11), the density-dependent term *N* increases by a factor of 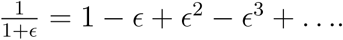 Thus, the deviations in *f* (*N)* are an order of magnitude smaller than those shown in Fig. (6).

Author contributions
JB and JM conceptualized the manuscript. JB did the formal analysis. JB wrote the manuscript with review and editing from JM

## References

P. Amarasekare. Interference competition and species coexistence. Proceedings of the Royal Society of London B: Biological Sciences, 269(1509):2541–2550, 2002.

N. Barton, D. Briggs, J. Eisen, D. Goldstein, and N. Patel. Evolution. NY: Cold Spring Harbor Laboratory Press, 2007.

M. Begon, J. L. Harper, and C. R. Townsend. Ecology. Individuals, populations and communities. 2nd edn. Blackwell scientific publications, 1990.

T. Benton and A. Grant. Evolutionary fitness in ecology: comparing measures of fitness in stochastic, density-dependent environments. Evolutionary Ecology Research, 2(6):769–789, 2000.

A. O. Bergland, E. L. Behrman, K. R. O’Brien, P. S. Schmidt, and D. A. Petrov. Genomic Evidence of Rapid and Stable Adaptive Oscillations over Seasonal Time Scales in Drosophila. PLOS Genetics, 10(11):1–19, 11 2014. doi: 10.1371/journal.pgen.1004775.

J. Bertram, K. Gomez, and J. Masel. Predicting patterns of long-term adaptation and extinction with population genetics. Evolution, 71(2):204–214, 2017.

B. M. Bolker and S. W. Pacala. Spatial moment equations for plant competition: Understanding spatial strategies and the advantages of short dispersal. The American Naturalist, 153(6):575–602, 1999. doi: 10.1086/303199.

M. S. Boyce. Restitution of r-and k-selection as a model of density-dependent natural selection. Annual Review of Ecology and Systematics, 15:427–447, 1984.

R. Burger and M. Lynch. Evolution and extinction in a changing environment: a quantitative-genetic analysis. Evolution, 49(1):151–163, 1995.

T. J. Case and M. E. Gilpin. Interference competition and niche theory. Proceedings of the National Academy of Sciences, 71(8):3073–3077, 1974.

B. Charlesworth. Selection in density-regulated populations. Ecology, 52(3):469–474, 1971.

B. Charlesworth. Evolution in age-structured populations, volume 2. Cambridge University Press Cambridge, 1994.

P. L. Chesson and R. R. Warner. Environmental variability promotes coexistence in lottery competitive systems. American Naturalist, 117(6):923–943, 1981.

T. Chotibut and D. R. Nelson. Population genetics with fluctuating population sizes. Journal of Statistical Physics, 167(3-4):777–791, 2017.

F. Christiansen. Density dependent selection. In Evolution of Population Biology: Modern Synthesis, pages 139–155. Cambridge University Press, 2004.

G. W. Constable and A. J. McKane. Mapping of the stochastic lotka-volterra model to models of population genetics and game theory. Physical Review E, 96(2):022416, 2017.

J. F. Crow, M. Kimura, et al. An introduction to population genetics theory. New York, Evanston and London: Harper & Row, Publishers, 1970.

U. Dieckmann and R. Ferrière. Adaptive dynamics and evolving biodiversity. 2004.

M. Doebeli, Y. Ispolatov, and B. Simon. Towards a mechanistic foundation of evolutionary theory. eLife, 6:e23804, Feb 2017. ISSN 2050-084X. doi: 10.7554/eLife.23804.

S. Engen, R. Lande, and B.-E. Saether. A quantitative genetic model of *r*- and *k*-selection in a fluctuating population. The American Naturalist, 181(6):725–736, 2013. ISSN 00030147, 15375323. URL http://www.jstor.org/stable/10.1086/670257.

W. J. Ewens. Mathematical Population Genetics 1: Theoretical Introduction. Springer Science & Business Media, 2004.

R. Ferríere and S. Legendre. Eco-evolutionary feedbacks, adaptive dynamics and evolutionary rescue theory. Phil. Trans. R. Soc. B, 368(1610):20120081, 2013.

R. A. Fisher. The genetical theory of natural selection: a complete variorum edition. Oxford University Press, 1930.

D. E. Gill. Intrinsic rate of increase, saturation density, and competitive ability. ii. the evolution of competitive ability. American Naturalist, 108:103–116, 1974.

J. H. Gillespie. Population genetics: a concise guide (2nd Ed.). John Hopkins University Press, 2010.

J. P. Grover. Resource competition, volume 19. Springer Science & Business Media, 1997.

J. B. S. Haldane. The cost of natural selection. Journal of Genetics, 55(3):511, 1957.

M. Kimura. Change of gene frequencies by natural selection under population number regulation. Proceedings of the National Academy of Sciences, 75(4):1934–1937, 1978.

M. Kimura and J. F. Crow. Natural selection and gene substitution. Genetics Research, 13 (2):127–141, 1969.

V. A. Kostitzin. Mathematical biology. George G. Harrap And Company Ltd.; London, 1939.

A. Lambert et al. The branching process with logistic growth. The Annals of Applied Probability, 15(2):1506–1535, 2005.

R. Lande, S. Engen, and B.-E. Sæther. An evolutionary maximum principle for density-dependent population dynamics in a fluctuating environment. Philosophical Transactions of the Royal Society B: Biological Sciences, 364(1523):1511–1518, 2009.

J. A. Leon and B. Charlesworth. Ecological versions of Fisher’s fundamental theorem of natural selection. Ecology, 59(3):457–464, 1978.

R. Levins and D. Culver. Regional coexistence of species and competition between rare species. Proceedings of the National Academy of Sciences, 68(6):1246–1248, 1971.

R. H. MacArthur. Some generalized theorems of natural selection. Proceedings of the National Academy of Sciences, 48(11):1893–1897, 1962.

R. H. MacArthur and E. O. Wilson. Theory of Island Biogeography. Princeton University Press, 1967.

J. Mallet. The struggle for existence. How the notion of carrying capacity, K, obscures the links between demography, Darwinian evolution and speciation. Evol Ecol Res, 14:627–665, 2012.

P. W. Messer, S. P. Ellner, and N. G. Hairston. Can population genetics adapt to rapid evolution? Trends in Genetics, 32(7):408–418, 2016.

C. J. E. Metcalf and S. Pavard. Why evolutionary biologists should be demographers. Trends in Ecology and Evolution, 22(4):205–212, 2007. ISSN 0169-5347. doi: https://doi.org/10.1016/j.tree.2006.12.001.

J. A. Metz, R. M. Nisbet, and S. A. Geritz. How should we define ‘fitness’ for general ecological scenarios? Trends in Ecology & Evolution, 7(6):198–202, 1992.

T. Nagylaki. Dynamics of density-and frequency-dependent selection. Proceedings of the National Academy of Sciences, 76(1):438–441, 1979.

T. Nagylaki et al. Introduction to theoretical population genetics, volume 142. Springer-Verlag Berlin, 1992.

M. Nei. Fertility excess necessary for gene substitution in regulated populations. Genetics, 68(1):169, 1971.

A. J. Nicholson. An outline of the dynamics of animal populations. Australian journal of Zoology, 2(1):9–65, 1954.

S. P. Otto and T. Day. A biologist’s guide to mathematical modeling in ecology and evolution. Princeton University Press, 2011.

T. L. Parsons and C. Quince. Fixation in haploid populations exhibiting density dependence i: the non-neutral case. Theoretical population biology, 72(1):121–135, 2007.

T. Prout. Some relationships between density-independent selection and density-dependent population growth. Evol. Biol, 13:1–68, 1980.

J. Roughgarden. Theory of population genetics and evolutionary ecology: an introduction. Macmillan New York NY United States, 1979.

P. F. Sale. Maintenance of high diversity in coral reef fish communities. The American Naturalist, 111(978):337–359, 1977.

P. E. Smouse. The implications of density-dependent population growth for frequency-and density-dependent selection. The American Naturalist, 110(975):849–860, 1976.

H. Svardal, C. Rueffler, and J. Hermisson. A general condition for adaptive genetic polymorphism in temporally and spatially heterogeneous environments. Theoretical Population Biology, 99:76–97, 2015. ISSN 0040-5809. doi: http://dx.doi.org/10.1016/j.tpb.2014.11.002.

D. Tilman. Competition and biodiversity in spatially structured habitats. Ecology, 75(1): 2–16, 1994.

J. Travis, J. Leips, and F. H. Rodd. Evolution in population parameters: Density-dependent selection or density-dependent fitness? The American Naturalist, 181(S1):S9–S20, 2013. doi: 10.1086/669970.

J. Turner and M. Williamson. Population size, natural selection and the genetic load. Nature, 218(5142):700–700, 1968.

G. P. Wagner. The measurement theory of fitness. Evolution, 64(5):1358–1376, 2010.

